# Therapeutic Treatment of Aerosolized Staphylococcal Enterotoxin B in Nonhuman Primates with two Monoclonal Antibodies

**DOI:** 10.1101/424036

**Authors:** Daniel Verreault, Jane Ennis, Kevin Whaley, Stephanie Z. Killeen, Hatice Karauzum, M.Javad Aman, Rick Holtsberg, Lara Doyle-Meyers, Peter J. Didier, Larry Zeitlin, Chad J. Roy

**Author notes:** Centre d’expertise en analyse environnementale du Québec, Québec Canada (Daniel Verreault). Corresponding Author: Dr. Chad J. Roy, TNPRC, Division of Microbiology, 18703 Three Rivers Road, Covington, LA 70433.

## Abstract

Staphylococcal enterotoxin B (SEB) is a protein exotoxin found on the cell surface of *Staphylococcus aureus* that is the source for multiple pathologies in man. When purified and concentrated in aerosol form, SEB can cause an acute and often fatal intoxication, and thus is considered a biological threat agent. There are currently no vaccines or treatments approved for human use. Studies in rodent models of SEB intoxication show that antibody therapy may be a promising treatment strategy, however many have used antibodies only prophylactically or well before any clinical signs of intoxication are apparent. We assessed and compared the protective efficacy of two monoclonal antibodies, Ig121 and c19F1, when administered after aerosol exposure in a uniformly lethal nonhuman primate model of SEB intoxication. Rhesus macaques were challenged using small particle aerosols of SEB, and then were infused intravenously with a single dose of either Ig121 or c19F1 (10 mg/kg) at either 0.5, 2 or 4 hours postexposure. Onset of clinical signs, hematological, and cytokine response in untreated controls confirmed the acute onset and potency of the toxin used in the challenge. All animals administered either Ig121 or c19F1 survived SEB challenge, whereas the untreated controls succumbed to SEB intoxication 30-48 hours postexposure. These results represent the successful therapeutic *in vivo* protection by two investigational drugs against SEB in a severe nonhuman primate disease model and punctuate the therapeutic value of monoclonal antibodies hold when faced with treatment options for SEB-induced toxicity in a postexposure setting.

One Sentence Summary: Two high-affinity monoclonal antibodies were tested for therapeutic efficacy using a rhesus macaque challenge model of aerosolized SEB

## Introduction

The staphylococcal enterotoxins (SEs) are a well-characterized family of proteins secreted by *Staphylococcus aureus* that known to be toxic at very low concentrations (1). Staphylococcal enterotoxin B (SEB) is a member of the family of superantigens (SAgs), which are microbial proteins that induce polyclonal T cell activation in contrast to conventional antigens that undergo proteolytic processing by antigen presenting cells (APCs) and are presented as a Major Histocompatibility Complex (MHC)/peptide complex (2-4). SAgs bypass these specific mechanisms of antigen presentation by binding outside the peptide binding groove of MHC class II on APCs and the variable region of T cell receptor (TCR) β chain on T cells (5-7). Cross-linking of MHC-II and TCR by SAgs activates both APCs and T cells. SAg binding activates 520% of circulating T cells bearing specific V beta regions, leading to massive release of pro-inflammatory cytokines, activation of cell-adhesion molecules, increased T-cell proliferation, and eventually T-cell apoptosis/anergy (8). This sequence of events can culminate in a life threatening condition clinically referred to as toxic shock syndrome (TSS) marked by cytokine storm, rash, hypotension, fever, multisystem dysfunction, and death (9).

SEB is a prototype SAg with a potential to be used as an airborne, foodborne or waterborne toxic agent and therefore classified by the CDC as select agent and by the US National Institutes of Health as a Category B priority pathogen. It was developed as a bioweapon in the 20th century due to its incapacitating or lethal nature at much lower dose than required by many chemical agents. SEB has been considered a high-risk toxin because of its relative ease of production, temperature-independent stability, and exquisite toxicity by the inhalation route. Inhalation of SEB aerosols far exceeds other modalities of exposure in terms of potency and deleterious effects, all initiating at a remarkably low (inhaled) dose. When inhaled, nanogram levels of SEB are incapacitating in human (ED_50_= 0.0004 μg/kg), while μg doses of SEB can be lethal (LD_50_ = 0.02 μg/kg) (1). Inhaled SEB initiates a nearly instantaneous response in the lung after inhalation, marked by neutrophilic influx, massive cytokine release, and marked pathological changes (10-14). Major osmotic shifts in the lung tissue from SEB inhalation results in a primarily localized inflammatory response which leads to progressive vascular leak, microcapillary hemorrhage, and alveolar flooding (10, 15, 16). The use of animal models of SEB intoxication to evaluate potential treatments are complicated by decreasing sensitivity based upon phylogenetic evolution; murine species are generally unresponsive to SEB unless genetically manipulated (17) or the reaction potentiated by coadministration of an agent such as LPS (15, 18). Nonhuman primate species have been shown to be the closest disease model to study pathophysiology of SEB-induced toxicity or in the testing of promising therapies and vaccine products (11, 19-22).

There are currently no vaccines or therapies approved by the U.S. Food and Drug Administration for either preventing or treating SEB intoxication by any modality of exposure. To date the development of a vaccine has been decidedly slow (23), although research has progressed on the development of STEBVax (24). Research and development on possible therapeutic agents has been even less successful. Known anti-inflammatories such as dexamethasone have been used with some success when administered prophylactically (25). Experimental treatments such as CD44 ligand, MyD88 mimetic, and IL-1 binding products have also shown promise (26-29) as treatment for SEB-induced shock and acute lung injury. Currently, the most promising post-exposure treatment for SEB intoxication appears to be from experimental monoclonal antibodies (mAb) engineered to target binding and effectively neutralizing any further processing of the molecule by host systems (30-33). The monoclonal antibody Ig121 was shown to protect against systemic exposure in a murine potentiation challenge model (34), although this particular effort did not include small particle aerosol exposure as a modality of challenge. The c19F1 mAB product has shown to be protective in follow-up murine SEB challenge studies in our laboratories, although the product was one of three mABs administered as a combination product (35).

In this study, we tested two monoclonal antibodies (IgG121, c19F1) in a severe model of aerosol SEB intoxication in the nonhuman primate. Each mAB targets different areas of the SEB molecule; Ig121 targets the T-cell receptor, whereas c19F1 binds to the variable β binding region of SEB (34). Binding differences between products are demonstrated *in vitro* using either SEB or the STEBVax vaccine product in preliminary experiments. In the animal experiments, we exposed naïve rhesus macaques to a lethal dose of aerosolized SEB, and treated via intravenous infusion at prescribed times after exposure (either 0.5, 2 or 4 hours) to test each mAB product to ameliorate and/or prevent the effects of intoxication.

## Results

### Binding ELISA

In order to evaluate the differences in potential binding regions between c19F1 and IgG121, ELISA assays were developed that use SEB or an attenuated form of SEB (STEBVax) as coating antigens. STEBVax is a recombinant mutated form of SEB containing three point mutations (L45R/Y89A/Y94A) that disrupt the interaction of the toxin with human MHC class II receptors and render the protein non-toxic while retaining the immunogenicity (36) . When these antibodies were tested in the two ELISAs, c19F1 was able to bind to SEB and STEBVax with a similar EC_50_. In contrast, IgG121 was able to bind to SEB but failed to bind to STEBVax at antibody concentrations as high as 1 μg/mL (Fig 1). These data suggest that IgG121 binds to the MHC binding surface of SEB while c19F1 binds to a distinct neutralizing site.

**Figure 1.**
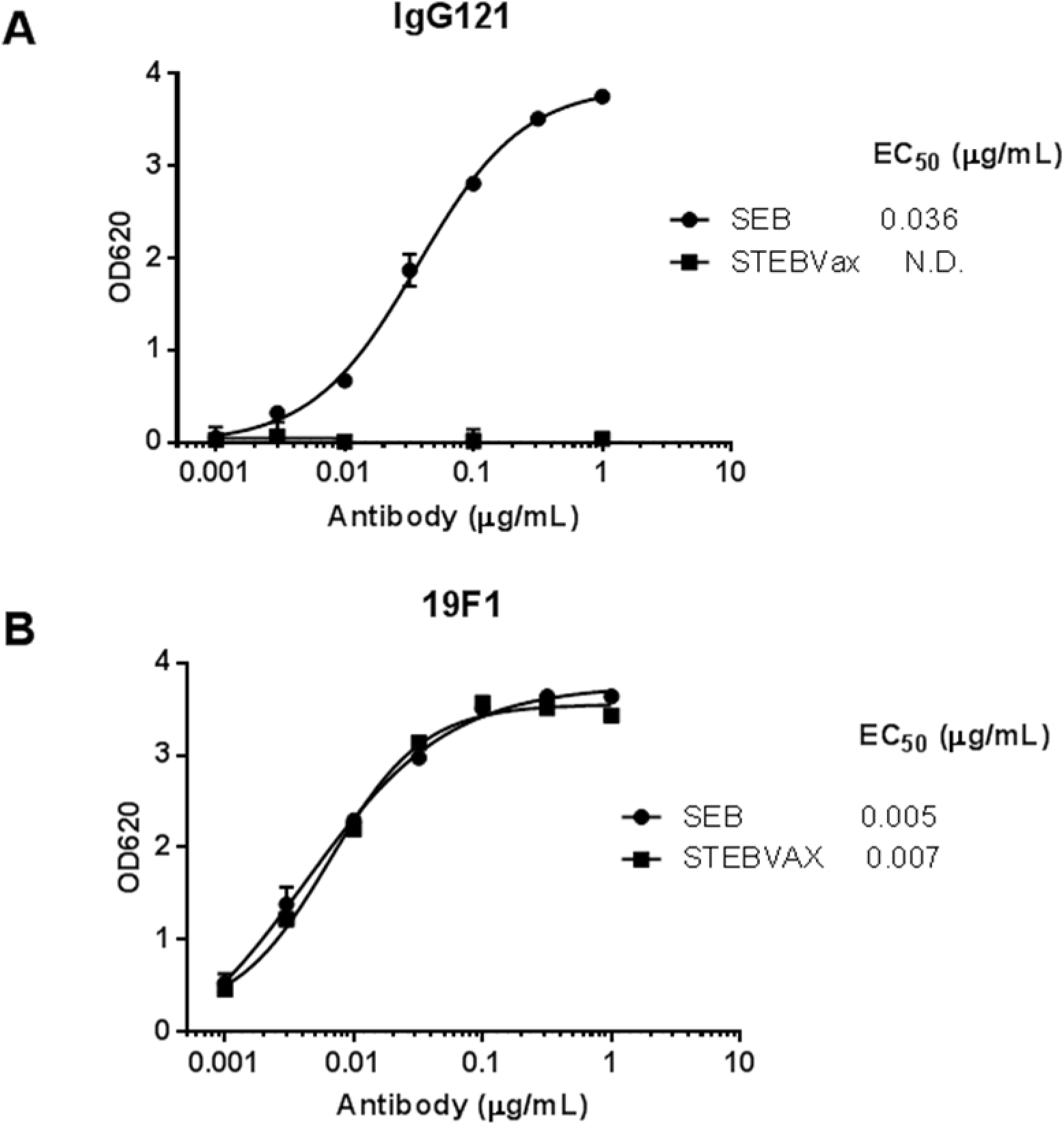
Binding of 121 and 19F1 to SEB or STEBVax. (A) Binding of 121 to SEB (closed circles) and STEBVax (Closed squares). (B) Binding of 19F1 to SEB (closed circles) and STEBVax (Closed squares).

### mAbs IgG121 and c19F1 protection of RM after SEB challenge

Rhesus macaques were challenged via the aerosol route with SEB toxin, and the challenge was measured as consistent between sham and antibody treated group (Fig. 2) The candidate mAbs were IV administered at a dose of 10 mg/kg at 0.5h or 4 hr (c19F1) or 2h or 4h (Ig121) and showed complete survival in mAb treated animals compared to zero survival in sham treated control groups (Fig 3).

**Figure 2.**
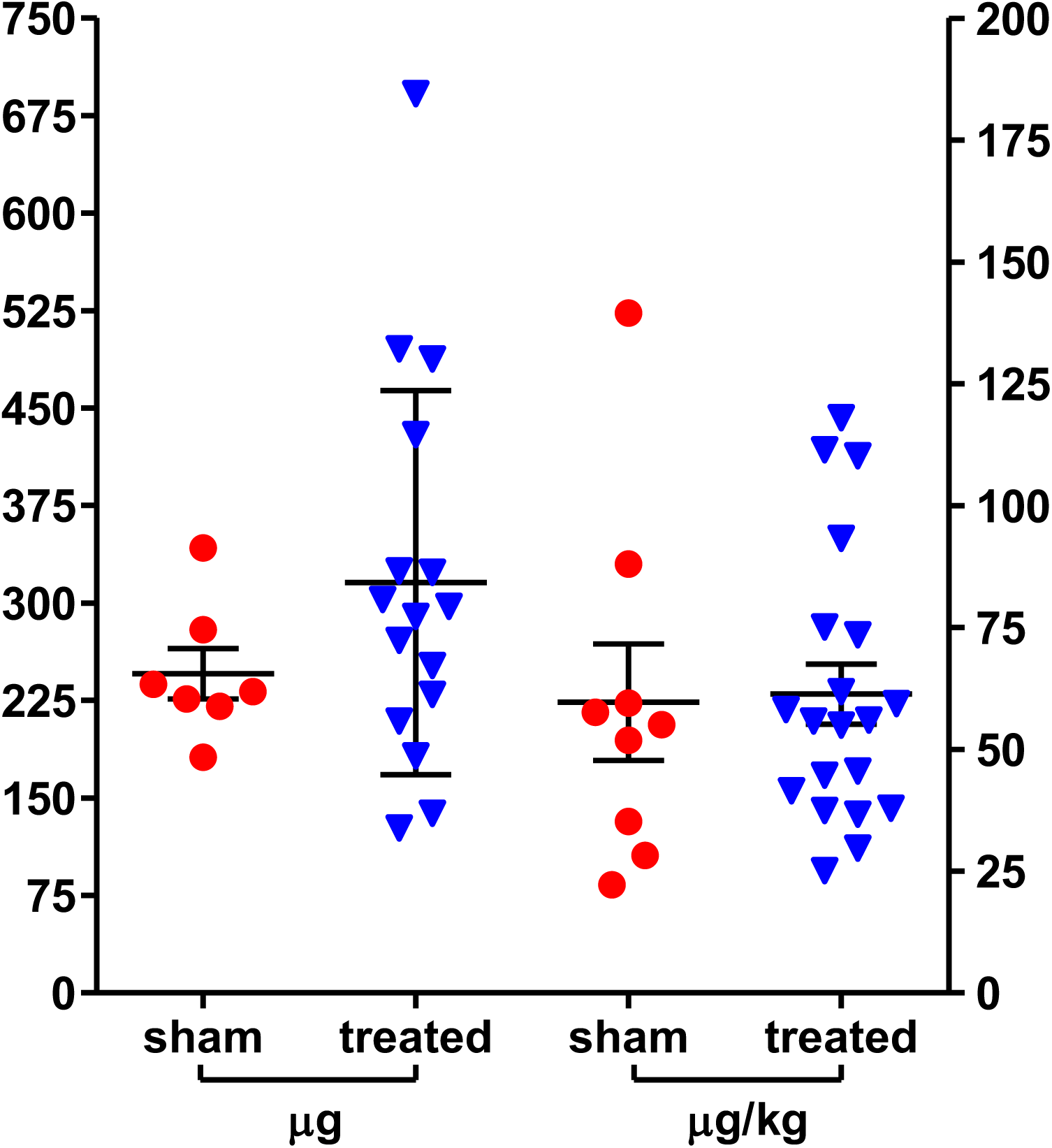
Individual inhaled challenge doses of rhesus macaques with aerosolized SEB toxin expressed as total μg inhaled and by a per weight basis (μg/kg). The line and error bars represent the mean and standard deviation for either antibody-treated or sham groups across all experiments for both measures (n=23). There were no statistically significant difference between challenge dose in group comparison of total μg (p=0.2392) or μg/kg values (p=0.8942).

**Figure 3.**
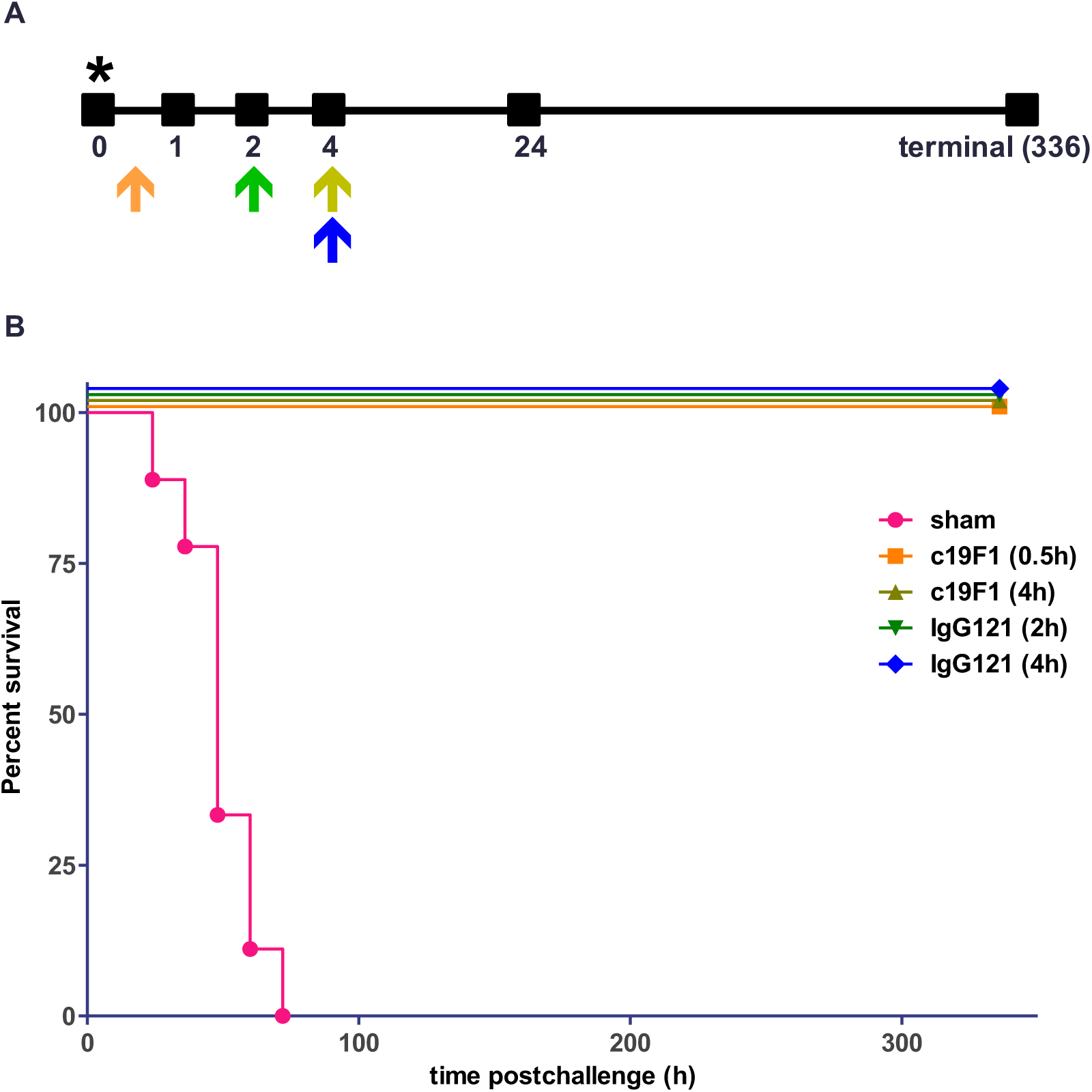
(A) Diagram of antibody treatment schedule, representing single dosage time point/group, and sampling time (hours) after SEB challenge in macaques. Orange arrow, c19F1 0.5 h treatment; olive arrow, c19F1 4 h treatment; green arrow, IgG121 2 h treatment; blue arrow, IgG121 4 hr treatment. Asterisk represents the hour of SEB challenge. (B) Kaplan-Meier survival curve for the sham group (red, n=9), c19F1 at 0.5h (orange, n=8), c19F1 at 4h (olive, n=4), IgG121 at 2h (green, n=4), and IgG121 at 4h (blue, n=4).

### Hematology

Hematological differences between control and treatment groups were minimal. mAb treated and sham groups showed a dramatic increase of neutrophils post-exposure when compared to pre-exposure values (Fig 4). Neutrophil percentages of values of the survivors in the treatment group were similar to that the control group animals. Similarly, lymphocyte changes in both groups were similar in profile; change in pre-exposure values significantly decreased post-exposure. Decreases were also observed in monocyte percentages.

**Figure 4.**
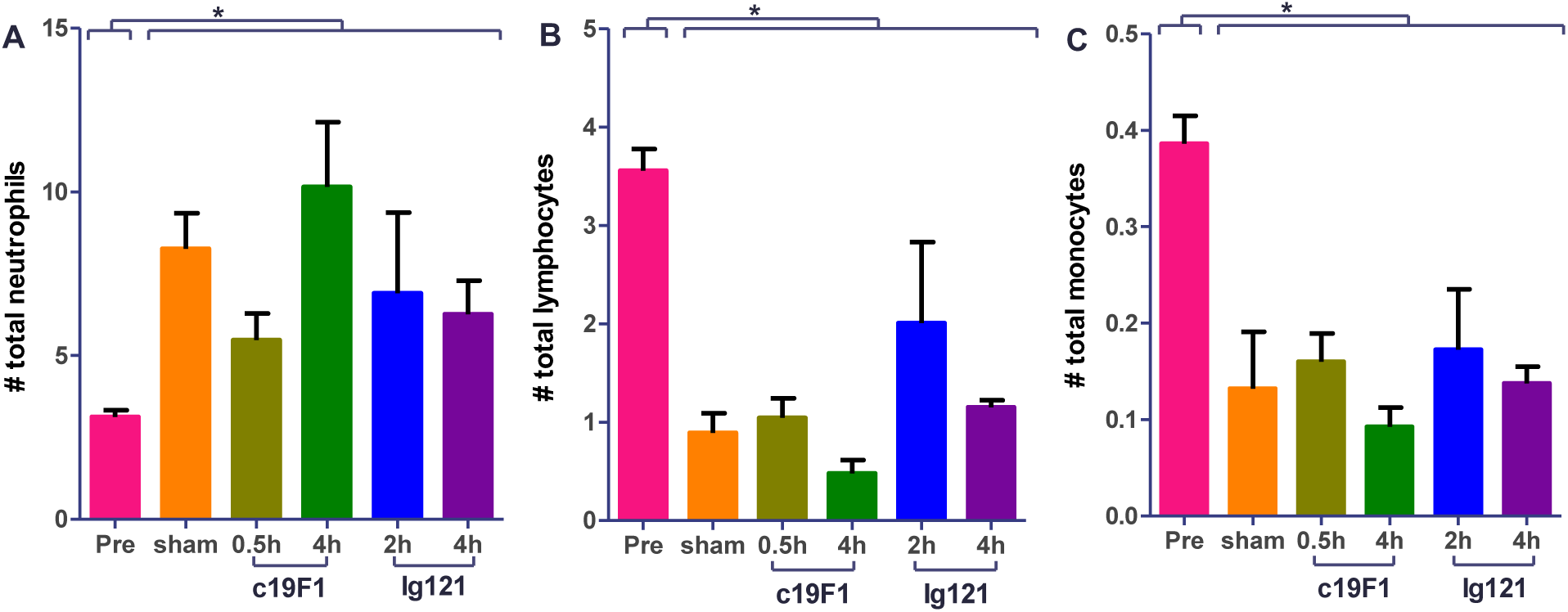
Cellularity from peripheral blood from all animals (n=23) prior to SEB challenge (Pre) and +24h post challenge and/or treatment with either antibody. Each graph represents A) neutrophils, B) lymphocytes, or C) monocyte count. All postchallenge values for either sham or treated animals were significantly different (^∗^ denotes p<0.05) when compared to preexposure values. There was no statistical difference between sham and treatment animals postexposure for any of the values presented.

### Clinical Chemistries

There were blood chemistry differences between the control group and treatment group. The average aspartate aminotransferase (AST) was observed to increase in sham control animals compared to pre-challenge levels (Fig 5), and the early time point mAb interventions for both c19F1 (0.5h) and IgG121 (2h) showed no change compared to pre-challenge control. The AST levels increased to levels comparable with sham control in the 4h mAb treated groups for both c19F1 and IgG121. Notable increase in alanine aminotransferase (ALT) was present in the sham control animals and the mAb-treated animals. Other parameters of note included blood urea nitrogen (BUN) and creatinine increases in all controls and all but 0.5h c19F1 (BUN) and 0.5h c19F1 and 4h IgG121 (creatinine) (Fig 6). Other notable changes were a significant decrease in serum protein in the 0.5h c19F1 treatment group and a reduction in serum albumin in sham control, and 4h c19F1 and 2h IgG121 treatment groups.

**Figure 5.**
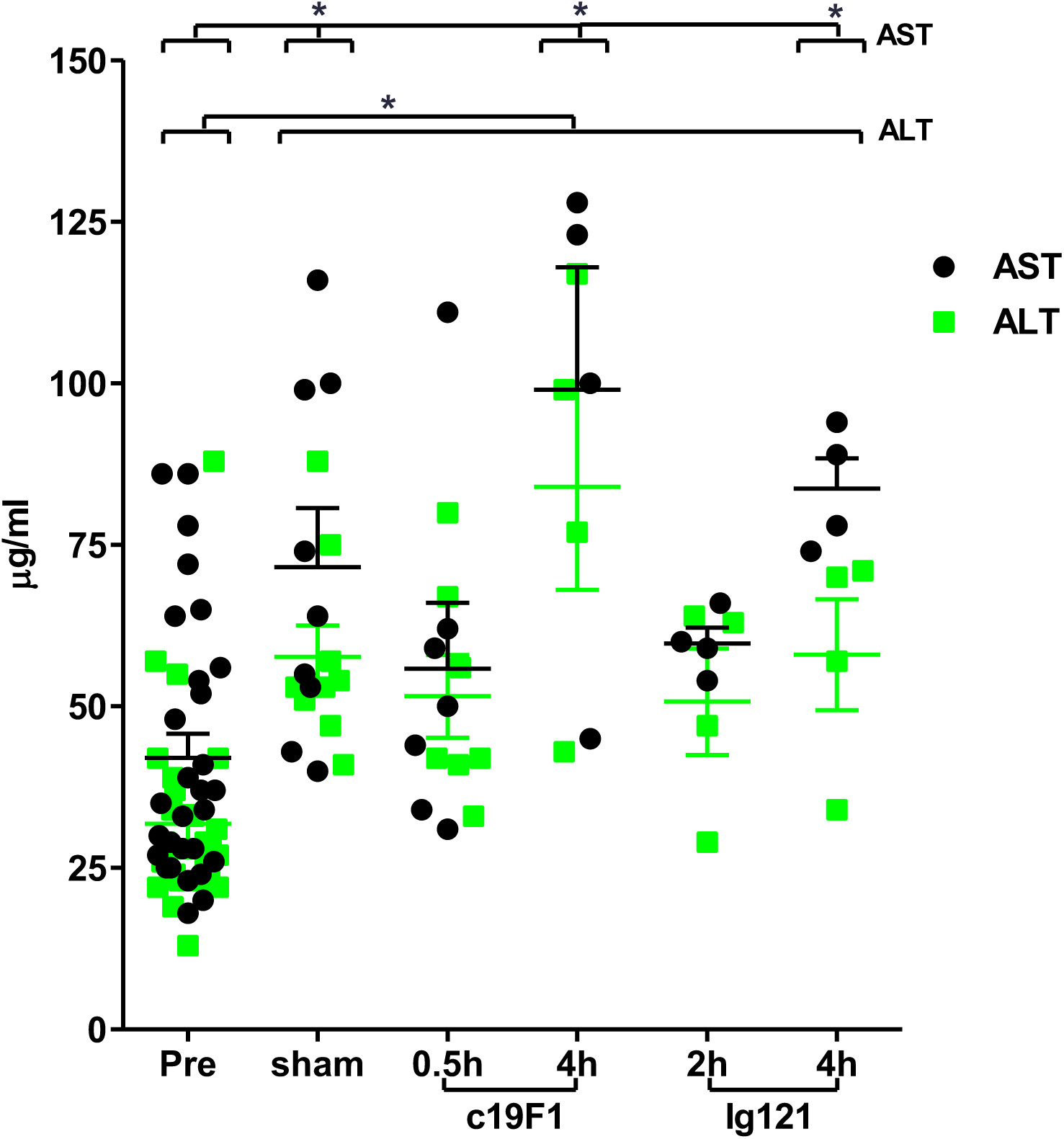
Aspartate aminotransferase (AST; closed circles) and alanine transaminase (ALT; squares) measured from peripheral blood from all animals (n=23) prior to SEB challenge (Pre) and +24h post challenge and/or treatment with either antibody. Significance differences for either sham or treated animals is denoted with ^∗^ (p<0.05) when compared to preexposure (Pre) values.

**Figure 6.**
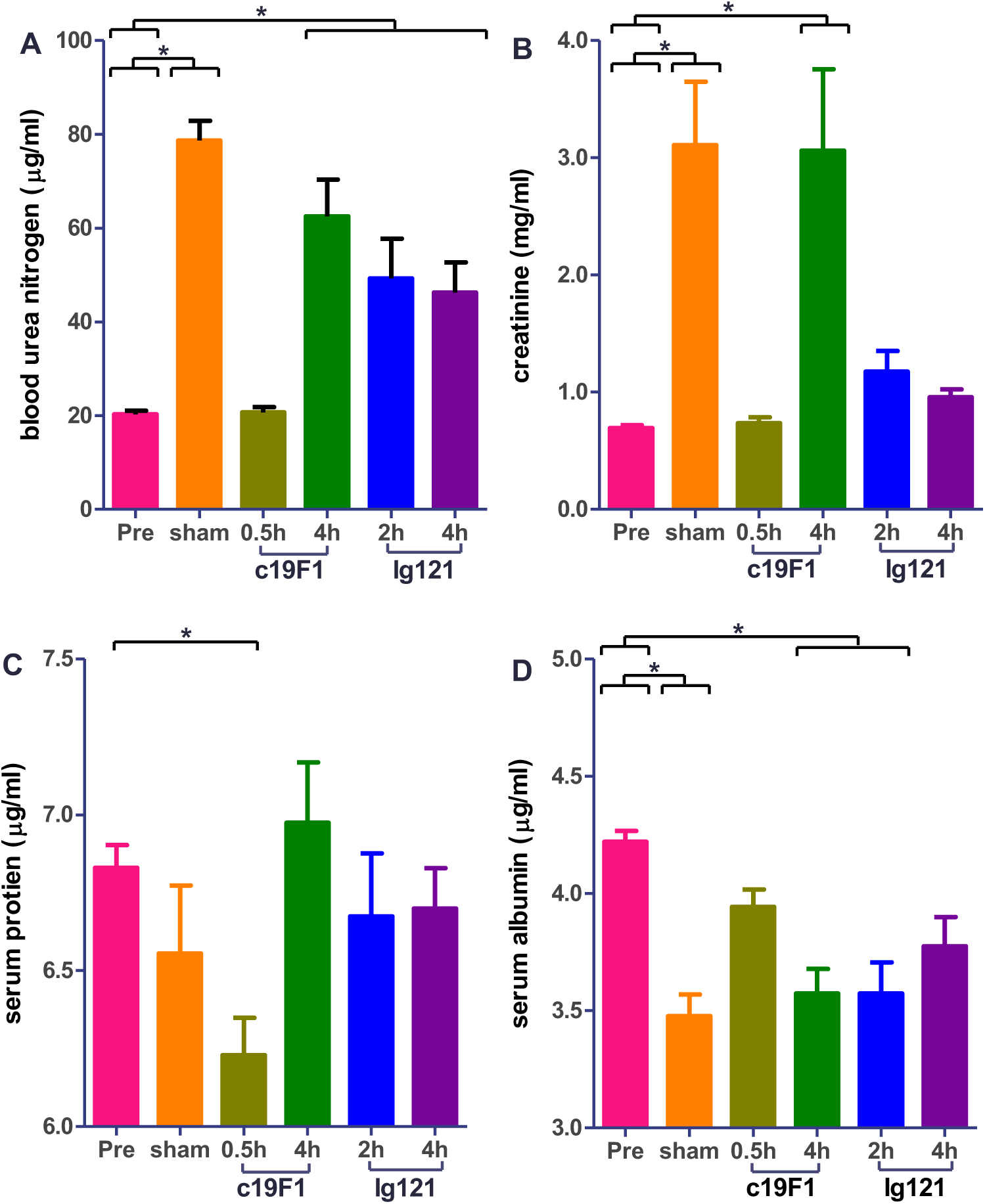
Relevant clinical chemistries derived from peripheral blood from all animals (n=23) prior to SEB challenge (Pre) and +24h post challenge and/or treatment with either antibody. Each graph shows A) blood urea nitrogen, B) creatinine, C) serum protein, and D) serum albumin. Significance differences for either sham or treated animals is denoted with ^∗^ (p<0.05) when compared to preexposure (Pre) values.

### Cytokines

Cytokine/chemokines were measured in replicate serum samples in all animals prior to and +24h and +48h post intoxication SEB challenge. Analysis was performed using a Bio-Plex^®^ 200 suspension array system, allowing analysis for 28 analytes in a single serum sample. Results of this analysis indicated a strong inflammatory systemic response as a consequence of SEB challenge, with discretion of cytokine response between treated and control animals (Fig 7). Notable lower concentrations were mainly early-phase proinflammatory cytokines and chemokines including interleukin-1 receptor antagonist (IL-1RA), macrophage migration inhibitory factor (MIF), monokine-induced gamma (MIG), monocyte attractant protein-1 (MCP1), eotaxin (eosinophilic chemoattractant), and interleukin-6 (IL-6) at the early (+0.5h) c19F1 treatment intervention at +24h PI when compared to preexposure levels. There was a noticeable distinction between the gradient of concentrations of cytokines and chemokines between timepoints in the course of response (+24h v. +48h) in both the treated and control animals.

**Figure 7.**
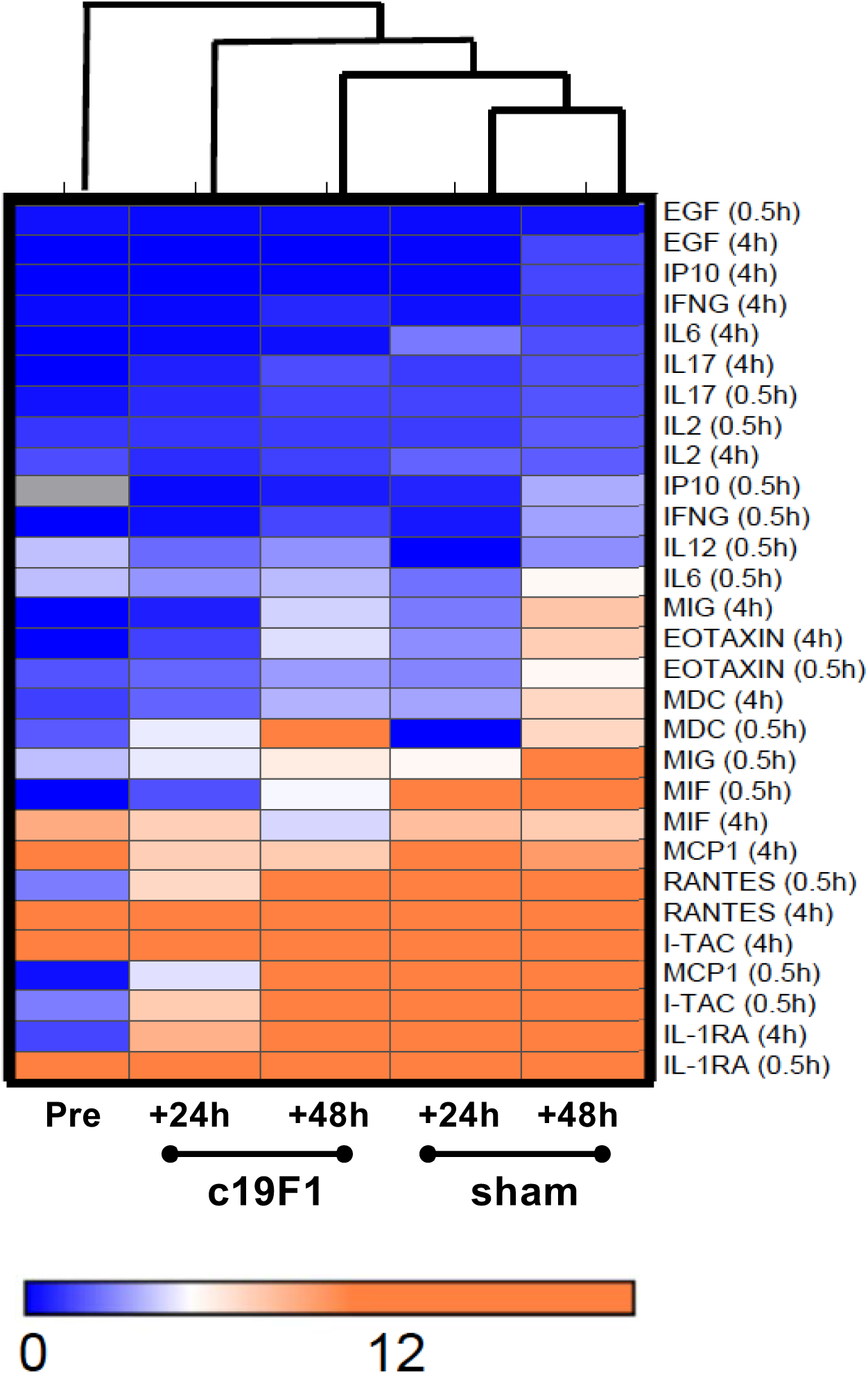
Hierarchical cluster analysis of cytokine data between c19F1-treated animals and controls. Each column represents blood collection timepoint prior to SEB exposure (Pre), or postexposure (either 24 or 48h) for c19F1 or sham-treated animal groups. Each horizontal category refers to time of treatment (either 0.5 or 4h PI) and corresponding timed sham treatments. Color legend is on the bottom of the figure. Orange indicates cytokines with a greater concentration relative to the geometrical mean of the Pre values, blue indicates cytokines with a lower concentration relative to the preexposure geometrical mean.

### Lung Pathology

Lung damage due to SEB aerosol exposure was measured through wet lung weight in all sham and mAb treatment groups. In all cases treatment with either c19F1 or IgG121 resulted in a significant reduction in wet lung weight, with no difference distinguishable among the treatment groups (Fig 8).

**Figure 8.**
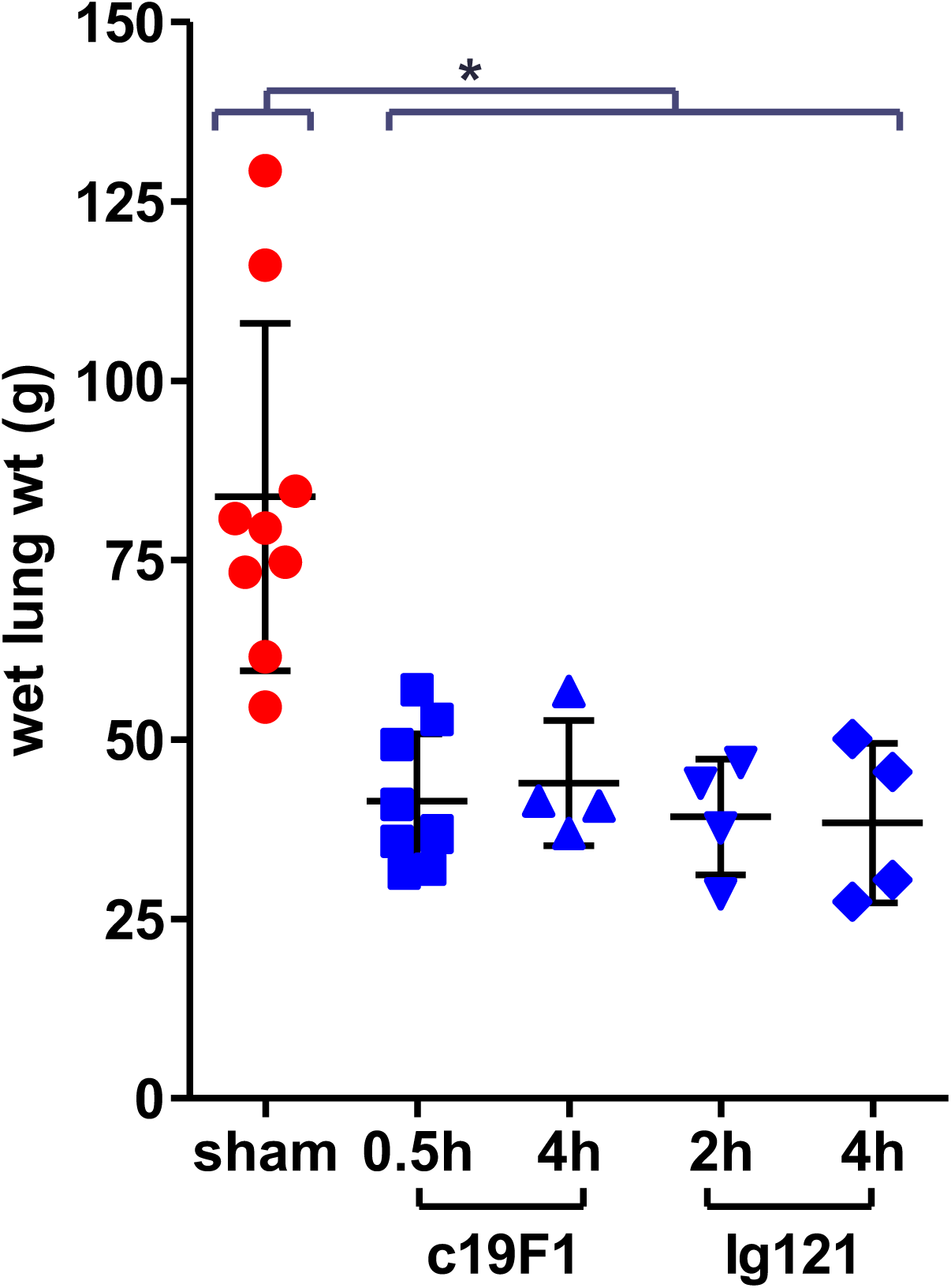
Wet lung weights of animals at necropsy; lines and error bars represent mean and standard deviation of each group. Weights of sham-treated animals were significantly different than all treated groups, ^∗^ denotes significance at (p<0.05). There was no significant difference in lung weight between treatment groups when compared.

### Histopathology

Five of six SEB-exposed sham treatment group animals developed severe pulmonary edema and mild intra-alveolar inflammation dominated by neutrophils with most demonstrating edema and acute mild inflammation in the bronchial node (Fig 9). One animal had minimal pulmonary edema without inflammation but acute randomly distributed necrotizing hepatitis and a small abscess in the mesenteric node consistent with bacterial infection. Mild chronic colitis was detected in four of six animals.

**Figure 9.**
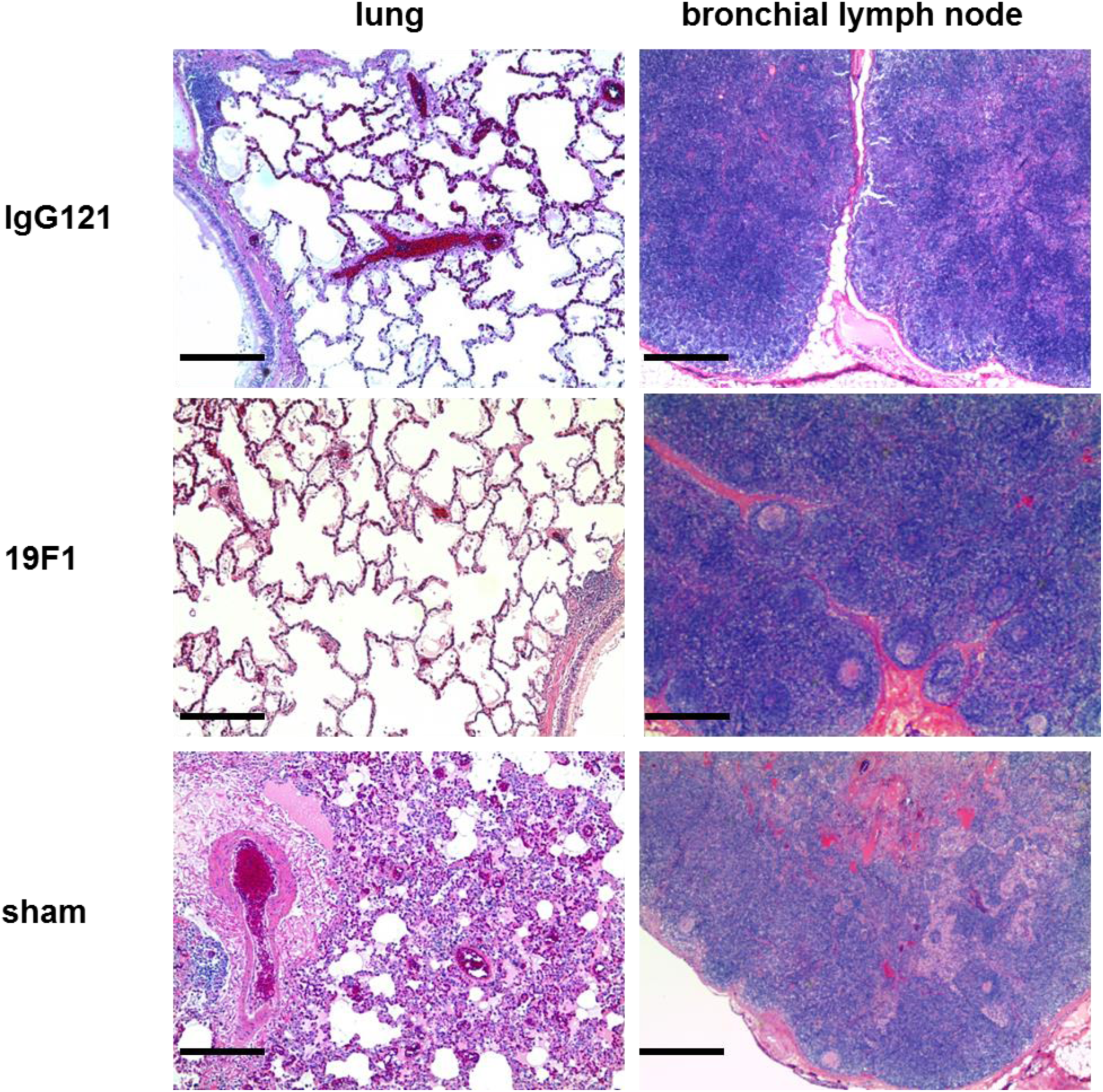
Representative tissues from mAb-treated (IgG121 and 19F1 at 10 mg/kg each) and sham untreated controls. Bar represents 100 μm.

SEB-exposed animals treated with IgG121 or c19F1 were nearly indistinguishable, as evidenced by similar categorical lesion scoring of lung tissue from both treatment groups in contrast to the sham-treated animals (Fig 10). Most had minimal to mild lymphoid hyperplasia in the lung, mild to moderate lymphoid hyperplasia of the bronchial node and spleen, and only three in the c19F1 group presented with minimal widely scattered foci of chronic inflammation in the lung. Edema was not present in lung or bronchial nodes, and only one animal in the c19F1 group had a minimal degree of colitis.

**Figure 10.**
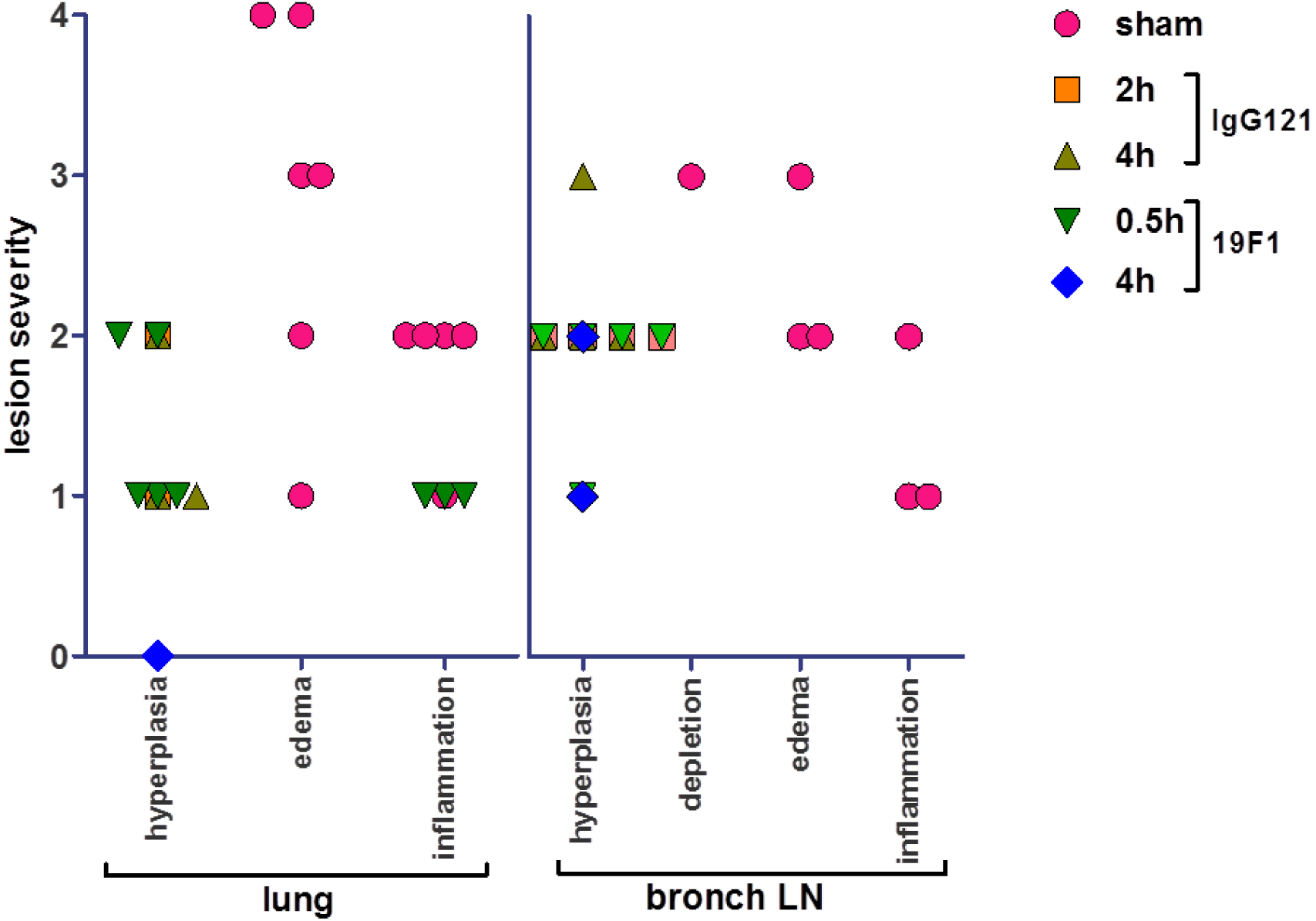
Categorical scoring of lesion severity in lungs and bronchial lymph nodes for major representative histological changes in each tissue type examined. Categories are represented as 0, normal; 1, minimal; 2, mild; 3, moderate; and 4, severe.

## Discussion

The misuse of biological toxins such as SEB remains a threat to both the military and civilian population. Recent attempts of poisonings through various mechanisms, including the U.S. postal service (37) is demonstrative of the ever-present threat and corresponding medical need for rapid acting countermeasures. In this study, we have addressed the possibility of providing efficacious post-exposure therapies to prevent the effects associated with SEB intoxication that would be suitable for use in humans after an accidental or intentional misuse exposure to SEB. The results reported are an important demonstration of a protective outcome in a nonhuman primate model of SEB intoxication and pathogenesis using one of two monoclonal antibodies. c19F1 and Ig121 are the only drug candidates demonstrated to be effective post-exposure against aerosolized SEB in the rhesus macaque model, which is uniformly lethal given the appropriate dose and modality of exposure (aerosol).

The animals that survived exposure through the therapeutic administration of either of the mAbs showed a different profile in clinical response than the untreated controls. Hematological and blood chemistry changes showed minimal differences when group responses were compared. Pathological outcome shows clear differences in damage in the control animals that died from exposure when compared to treated survivors which showed minimal changes in the lung as a result of aerosol exposure. Together, the data derived from the efficacy experiments to date indicate that the anti-SEB mAbs c19F1 and IgG121 are the only effective antitoxin therapies that have been identified that could potentially be used to treat intoxication in humans. In order to develop these mAbs as treatments for SEB intoxication however, further work is needed. Elucidation of the mechanism of action, binding sites, and toxicological studies are needed to ensure there are no safety concerns. In addition, the prospect of humanizing c19F1 in order to increase protection while reducing unwanted human immunogenicity may be warranted.

## Materials and Methods

### Study Design

An initial pharmacokinetics study was performed at TNPRC to determine the optimal route of administration for each of the antibodies (data not shown) where animals were administered either antibody either IM or IV and bled according to a predetermined schedule. Serum was analyzed via Biacore (L. Zeitlin) for determination of concentration of the antibody. Intravenous infusion was determined the optimal route of administration based upon the rapidity (<30 min) which the antibodies reached serum levels consistent with protection. In the therapeutics studies, all animals were first exposed to SEB aerosols at a target dose consistent with lethality. Thereafter groups were administered either c19F1 or IgG1 via IV infusion at equivalent dose (10 mg/kg) at prescribed times (0.5, 2, or 4h) postexposure. Sham treated animals were administered sterile water for injection by that same route.

### Animals

Age- and sex-matched rhesus macaques (*Macaca mulatta*) weighing between 4-8 kg, free of simian immunodeficiency virus (SIV), simian type D retrovirus, simian T-lymphotropic virus were used. All experiments using macaques were approved by the Tulane Institutional Animal Care and Use Committee. The Tulane National Primate Research Center (TNPRC) is an Association for Assessment and Accreditation of Laboratory Animal Care International accredited facility (AAALAC#000594). The U.S. National Institutes of Health (NIH) Office of Laboratory Animal Welfare assurance number for the TNPRC is A3071-01. Nonhuman primate housing consisted of individual open metal caging units that allowed visual recognition and protected contact with other study animals in the room. Animals were maintained on standard primate chow supplemented daily with fresh fruits and vegetables. Animals were provided standard environmental enrichment during this study, which included manipulable items in cage, perches, foraging/task-oriented feeding methods, and human interactions with caretakers and research staff. All clinical procedures, including administration of anesthesia and analgesics, were carried out under the direction of a laboratory animal veterinarian. Animals were anesthetized with ketamine hydrochloride for blood collection procedures. All possible measures are taken to minimize discomfort of all the animals used in this study. Animals were closely monitored daily following surgery for any signs of illness such as anorexia, lethargy, diarrhea, vomiting, and dehydration. Appropriate medical care was implemented if any of these signs of illness were noted. If euthanasia was required in the judgment of the TNPRC veterinary staff, animals were euthanized in accordance with the recommendations of the panel on Euthanasia of the American Veterinary Medical Association. The standard method of euthanasia for nonhuman primates at the TNPRC is anesthesia with ketamine hydrochloride (10 mg/kg) followed by an overdose of sodium pentobarbital. Tulane University complies with NIH policy on animal welfare, the Animal Welfare Act, and all other applicable federal, state and local laws.

### SEB binding ELISA

Enzyme-linked immunosorbent assay (ELISA). Purified SEB or attenuated SEB (STEBVax) were immobilized at 200 ng/well on 96-well Nunc MaxiSorp plates (ThermoFisher Scientific) and incubated with serial dilutions of purified antibodies. Bound antibodies were detected using an HRP-conjugated anti-human secondary antibody (KPL) and TMB substrate (Life Technologies). Absorbance values determined at 620 nm were transformed using Softmax^®^ 4 parameter curve-fit (Molecular Devices). Half maximal effective concentration (EC_50_) value at the inflection point of the curve was determined.

### Animal Challenge and Therapy

SEB toxin was obtained from BEI resources (Manassas, VA) and was accompanied by results of purity testing shown by electrophoresis. The toxin was reconstituted in phosphate buffered saline immediately before being placed into the aerosol generating nebulizer. Animals were exposed to the toxin using a dynamic head-only inhalation exposure apparatus controlled by an electronic flow process platform (Biaera Technologies, Hagerstown, MD) and has been described previously (38, 39). Briefly, anesthetized animals are transported into a class III biological safety cabinet outfitted with the head-only inhalation configuration. The aerosol concentration and the estimated inhaled dose were calculated as described (40) using the minute volume measured during plethsymography and the SEB concentration in the aerosol as determined by protein assays of the all-glass impinger samples.

### Antibodies

The antibodies were provided by IBT (IgG121) and Mapp (c19F1) as a liquid product in a sealed glass vacuum vial. The doses for the evaluation were derived from the starting concentration and administration of the antibody was based upon prevailing body weights of the animals enrolled in the study. During the time between the extracting and use of the antibody, the sterile syringes were stored at 4 degrees C (in the dark) until administration. Care was taken to ensure that the antibody remained in solution prior to administration.

### Hematology, serum biochemistry, and blood coagulation

Hematologic analysis was performed with blood samples collected in EDTA anticoagulant using a Sysmex XT-2000i Analyzer and analysis of serum from clotted samples of blood was done on an Olympus AU400 chemistry analyzer with results automatically downloaded to the Animal Record System.

### Cytokine Analysis

Cytokine levels in serum were assayed using the Milliplex^®^ MAP Primate/Rhesus Cytokine/Chemokine Polystyrene Bead Panel (PRCYTOMAG-40K, Millipore Corp. (Billerica, MA)). This assay provides analysis for selected cytokines/chemokines within a single sample, which are G-CSF, VEGF, TNF-α, TGF-α, sCD40L, MIP-1β, MIP-1α, MCP-1, IL-18, IL-17, IL-15, IL-13, IL-12/23 (p40), IL-10, IL-8, IL-6, IL-5, IL-4, IL-2, IL-1β, IL-1rα, IFN-γ, and GM-CSF. Briefly, collected blood serum samples from the primates are disturbed by vortex, and then individually clarified through filter spin columns (catalog no. UFC30DV00, Millipore Corp.) by spinning at 12,000 × g for 4 min at room temperature. Each standard, control, or undiluted sample, in 25 μl, is added in duplicate to antibody-conjugated beads and incubated in a 96-well filter plate overnight at 2–8 °C with shaking at 650 rpm. After 16–18 h, wells are washed, and 25 μl of detection antibody is added to each well. After 1 h of incubating at room temperature with shaking, 25 μl of streptavidin-phycoerythrin is added to each well and incubated for 30 min with shaking. After final washes are completed, 150 μl of sheath fluid is added to each well. The plate is analyzed using a Bio-Plex^®^ 200 suspension array system (Bio-Rad). The instrument settings are as follows: 50 events/bead, 100-μl sample size, and gate settings at 8000–15,000. The software used to perform the assay and analyze data is Bio-Plex ManagerTM version 6.0, which calculates concentrations in pg/ml based on the respective standard curve for each cytokine.

### Histopathology

Tissue samples were fixed for 48 hr in zinc-modified formalin (Z-Fix, Anatech, LTD, Battle Creek, MI), dehydrated in a series of alcohols, embedded in paraffin, and sectioned at 5 μm thickness, before deparaffinized sections were stained with hematoxylin and eosin. Sections were examined by light microscopy on a Leica DMLB microscope equipped with a Leica EC3 camera.

### Statistical Analysis

Data were analyzed using PRISM software (GraphPad Software, Inc.). *In vivo* survival curves were analyzed using the log-rank (Mantel Cox) test.

## Abbreviations used

ALT: alanine aminotransferase
APCs: antigen presenting cells
AST: aspartate aminotransferase
AAALAC: Association for the Assessment and Accreditation of Laboratory Animal Care International
BUN: blood urea nitrogen
CDC: Centers for Disease Prevention and Control
ED_50_: effective dose 50%
ELISA: enzyme linked immunosorbent assay
G-CSF: granulocyte colony-stimulating factor
GM-CSF: granulocyte macrophage colony stimulating factor
HRP: horseradish perioxidase
IL-18: interleukin 18
IL-17: interleukin 17
IL-15: interleukin 15
IL-13: interleukin 13
IL-12/23: interleukin12/23
IL-10: interleukin 10
IL-8: interleukin 8
IL-6: interleukin 6
IL-5: interleukin 5
IL-4: interleukin 4
IL-2: interleukin 2
IL-1β: interleukin 1 beta
IL-1ra: interleukin 1 receptor antagonist
IM: intramuscular
IV: intravenous
IFN-γ: interferon gamma
LD_50_: lethal dose 50%
LPS: lipopolysaccharide
MHC: major histocompatibility complex
MIP-1β: macrophage inflammatory protein-1 beta
MIP-1α: macrophage inflammatory protein-1 alpha
MCP-1: monocyte chemoattractant protein-1
NIH: National Institutes of Health
SIV: simian immunodeficiency virus
sCD40L: soluble CD40-ligand
SEB: staphylococcal enterotoxin B
SAgs: superantigens
TCR: T cell receptor
TSS: toxic shock syndrome
TMB: 3,3’,5,5’-Tetramethylbenzidine
TNPRC: Tulane National Primate Research Center
TNF-α: tumor necrosis factor-alpha
TGF-α: tumor granulocyte factor-alpha
VEGF: vascular endothelial growth factor

